# Genomic and transcriptomic insights into the regulation of snake venom production

**DOI:** 10.1101/008474

**Authors:** Adam D. Hargreaves, Martin T. Swain, Matthew J. Hegarty, Darren W. Logan, John F. Mulley

## Abstract

The gene regulatory mechanisms underlying the rapid replenishment of snake venom following expenditure are currently unknown. Using a comparative transcriptomic approach we find that venomous and non-venomous species produce similar numbers of secreted products in their venom or salivary glands and that only one transcription factor (*Tbx3*) is expressed in venom glands but not salivary glands. We also find evidence for temporal variation in venom production. We have generated a draft genome sequence for the painted saw-scaled viper, *Echis coloratus*, and identified conserved transcription factor binding sites in the upstream regions of venom genes. We find binding sites to be conserved across members of the same gene family, but not between gene families, indicating that multiple gene regulatory networks are involved in venom production. Finally, we suggest that negative regulation may be important for rapid activation of the venom replenishment cycle.

## INTRODUCTION

Bites from venomous snakes, with their associated fatalities and long term physical and psychological effects, are an important public health issue in many parts of the developing world, often exacerbated by poor availability of health care and particularly a lack of effective and affordable antivenom treatments (Harrison et al. 2009; Mohapatra et al. 2011; Warrell 2010). It has been estimated that between 1.2-5.5 million snakebites occur annually, resulting in almost half a million envenomings and 20,000-90,000 deaths (Kasturiratne et al. 2008). Whilst there is obviously a need to address these issues through improvements in the quality and availability of healthcare in the rural tropics, there is also a requirement for research into the molecular basis of inter-and intra-specific variation in venom composition, which has implications for antivenom efficacy (Fry et al. 2003; Gutiérrez et al. 2009; Sunagar et al. 2014). Such variation can be a product of both genome-level and gene-level events, reflecting either variation in gene presence/absence between individuals and populations (as has been demonstrated for the presence or absence of Mojave toxin in the Mojave rattlesnake *Crotalus scutulatus scutulatus* (Wooldridge et al. 2001)) or changes in gene regulation that alter temporal, spatial or quantitative expression of the target genes (for example ontogenetic changes in venom composition in the South American pit vipers *Bothrops asper* (Alape-Girón et al. 2008), *Bothrops insularis* (Zelanis et al. 2007; Zelanis et al. 2008) and *Bothrops atrox* (Guércio et al. 2006; López-Lozano et al. 2002)).

Understanding the genetic basis of the regulation of snake venom production can inform our understanding of general evolutionary processes, such as the origin and fate of duplicate genes and the molecular-level mechanisms underpinning evolutionary innovation. Snake venom can be considered to represent such an innovation and has been hypothesised to diversify via gene duplication (Kordiš and Gubenšek 2000; Wong and Belov 2012), where novel toxins are “recruited” into the venom gland following duplication of a gene with a role elsewhere in the body and recruitment of one of the copies into the venom gland where selection can act to develop or increase toxicity (Casewell et al. 2013; Fry 2005; Lynch 2007). In order for genes to gain a new site of expression (neofunctionalisation (Force et al. 1999)) they must evolve new combinations of transcription factor binding sites in their associated *cis*-regulatory regions. We have recently shown that snake venom in fact evolves via the duplication and restriction of genes, rather than recruitment (Hargreaves et al. 2014a), and this process of subfunctionalisation rather than neofunctionalisation does not require the *de novo* creation of novel transcription factor binding sites.

Following venom expenditure (either through envenomation of prey or induced manually by “milking”) the production of toxic venom components must be initiated or upregulated and, given the energetic costs associated with venom production (McCue 2006) it must be assumed that this will be an efficient and rapid process (although these costs may not be as high as those related to other physiological processes such as digestion and shedding (Pintor et al. 2010)). Historically, a window 3-4 days post-milking has been taken to represent a peak period of venom synthesis (Boldrini-França et al. 2009; Kochva 1978; Paine et al. 1992; Wagstaff and Harrison 2006), although there has long been evidence to suggest that some venom components are transcribed almost immediately following (and certainly within a few hours of) venom expenditure (Currier et al. 2012; Lachumanan et al. 1999). Many previous studies of snake venom composition have understandably focussed on the period of peak synthesis (see for example, (Boldrini-França et al. 2009; Casewell et al. 2009; Margres et al. 2013; Neiva et al. 2009; Wagstaff and Harrison 2006)) and are therefore unable to provide insights into the signalling pathways and transcription factors that may regulate the production of snake venom. Whilst Ca^2+^ mobilisation, α-and β-adrenoceptors, the MAP kinase (ERK1/2) signalling pathway and nuclear factor κB (NFκB) and activator protein 1 (AP-1) transcription factors have been implicated in the initiation of venom production following milking in the South American pit viper *Bothrops jararaca* (Kerchove et al. 2008; Kerchove et al. 2004; Luna et al. 2009; Yamanouye et al. 2000), it is not known how widespread these putative gene regulatory network components might be, nor what other transcription factors and signalling pathways might be involved in the control of snake venom production. In addition, little is currently known about promoter and enhancer regions associated with venom genes, although binding sites for NFκB, *Sp1* and *Nf1* transcription factors have been identified upstream of the crotamine gene in the South American rattlesnake, *Crotalus durissus terrificus* (Rádis-Baptista et al. 2003). Binding sites for some of the proposed transcription factors have also been found in the promoter regions of three-finger toxins present in *Boiga dendrophilia* (*Nf1*) (Pawlak and Kini 2008), *Naja sputatrix* (NFκB, AP-1, *Nf1*, *Sp1*) (Lachumanan et al. 1998; Ma et al. 2001) and *Naja atra* (*Nf1*, *Sp1*) (Chang et al. 2004); and in Phospholipase A_2_ group IB genes of *Bungarus multicinctus* (*Sp1*) (Chu and Chang 2002)and *N. sputatrix* (AP-1, *Sp1*) (Jeyaseelan et al. 2000). It is worth noting that, to our knowledge, no studies of the promoter regions of venom genes in true vipers (Viperinae) are currently available in the literature.

We set out to determine the transcription factors and signalling pathways involved in the regulation of venom production through the use of comparative genomic and transcriptomic approaches in a range of reptile species. Our findings include the following: out of a set of seven body tissues from the painted saw-scaled viper (*Echis coloratus*) the venom gland expresses the lowest number of unique transcripts; the transcription factor and signalling pathway components of the venom glands of venomous species and the salivary glands of non-venomous species are highly similar; there is a shift of focus between transcription and translation during the venom replenishment cycle; and the expression of venom toxins may show temporal differences.

Finally, our analysis of the genomic regions upstream of venom genes reveals conserved transcription factor binding sites between members of the same gene family, but not between different gene families, suggesting a role for multiple gene regulatory networks in the production of snake venom. The close proximity of these binding sites to the start of venom genes is an arrangement likely to facilitate the duplication and diversification of toxin gene families whilst retaining required *cis*-regulatory regions.

## RESULTS

We have conducted the first large-scale comparative analysis of the regulation of snake venom production, utilising a transcriptomic analysis of the salivary or venom glands of six species of reptile (Figure 1), including two venomous vipers (the painted saw-scaled viper *Echis coloratus* and the Egyptian saw-scaled viper *Echis pyramidum)*, two non-venomous colubrids (the corn snake *Pantherophis guttatus* and rough green snake *Opheodrys aestivus*), a non-venomous boid (the royal python *Python regius*) and a member of the most basal lineage of squamates (the leopard gecko *Eublepharis macularius*). Six additional body tissues (brain, kidney, liver, ovary, cloacal scent gland and skin) were also sequenced from the painted saw-scaled viper and we have taken advantage of recently published venom gland and body tissue transcriptome data from the king cobra (Vonk et al. 2013) (*Ophiophagus hannah*), and venom gland transcriptomes for the Eastern diamondback rattlesnake (Rokyta et al. 2012) (*Crotalus adamanteus*) and Eastern coral snake (Margres et al. 2013) (*Micrurus fulvius*). We have also generated a low coverage draft whole genome assembly for the painted saw-scaled viper and have carried out comparative genomic analyses utilising previously published venom gene sequences and whole genome sequences for the king cobra (Vonk et al. 2013), Burmese python (Castoe et al. 2013) (*Python molurus bivittatus*) and Boa constrictor (Bradnam et al. 2013) (*Boa constrictor constrictor*).

**Figure 1.**
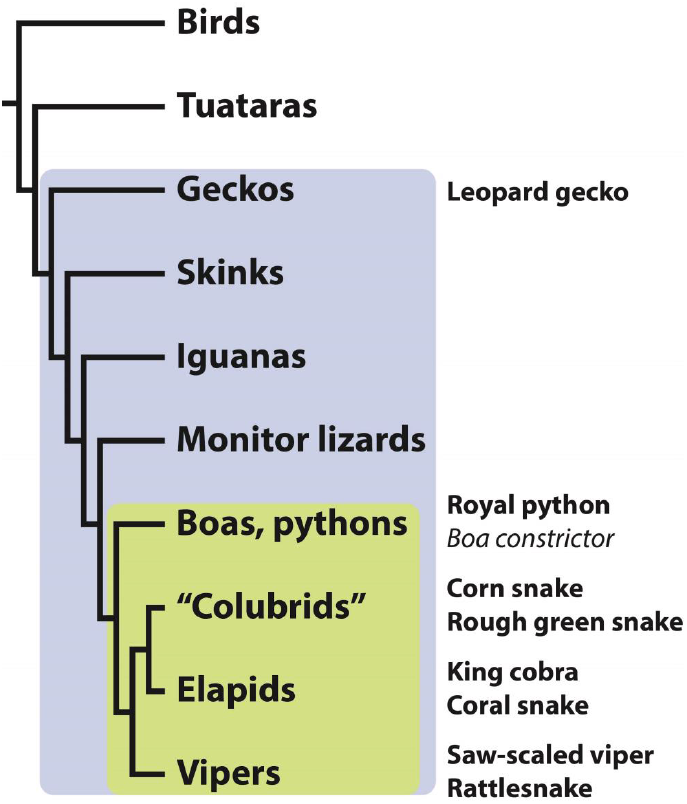
Simplified reptile phylogeny showing relationships of the study species. Squamate reptiles are shaded blue and snakes are green.

### Transcriptomics

Assembled transcriptomes contained between 18,169 and 134,236 contigs of ≥300bp and contig N50 values ranged from 950bp to 3,552bp (full assembly statistics are available in Supplementary table S1). Most contigs encoded an ORF of at least 20aa and of these between 3.72% and 10.64% were predicted to encode a signal peptide and so are likely to be secreted (Supplementary table S2-S4 and next section). Gene ontology was assigned to contigs using BLAST2GO (Conesa et al. 2005) and in order to provide a broad overview of the assigned gene ontologies we generated generic GOSlim annotations. Venom and salivary gland compositions appear to be broadly similar and comparisons of the venom gland and other body tissues in *E. coloratus* do not highlight any major differences (Supplementary figures S1-S6). A global *E. coloratus* tissue assembly containing 147,966 contigs of ≥300bp was created and the tissue distribution of these transcripts was determined by mapping sequencing reads from each tissue to this assembly. We identified 11,570 transcripts common to all 7 tissues (Figure 2) and suggest that these transcripts most likely represent ubiquitous maintenance or housekeeping genes common to all cells. Just 2,965 transcripts were found to be uniquely expressed in the venom gland, accounting for 8.27% of the total transcripts expressed in this tissue (far fewer than any other body tissues (Figure 2, Supplementary table S5)), although it may be possible that highly expressed venom genes are drowning out some lowly-expressed transcripts. Within these venom gland-specific transcripts we find a number that encode members of known venom toxin gene families, including three contigs encoding group III SVMPs, eight contigs for both metalloproteinases and serine protease and four contigs encoding C-type lectins (see Supplementary figures S7-S9 for GO annotation graphs of these unique transcripts). Interestingly, the venom gland shares 26,181 transcripts in common with the scent gland, more than with any other body tissue (Supplementary table S6) and this may reflect their similar functions as secretory tissues.

**Figure 2.**
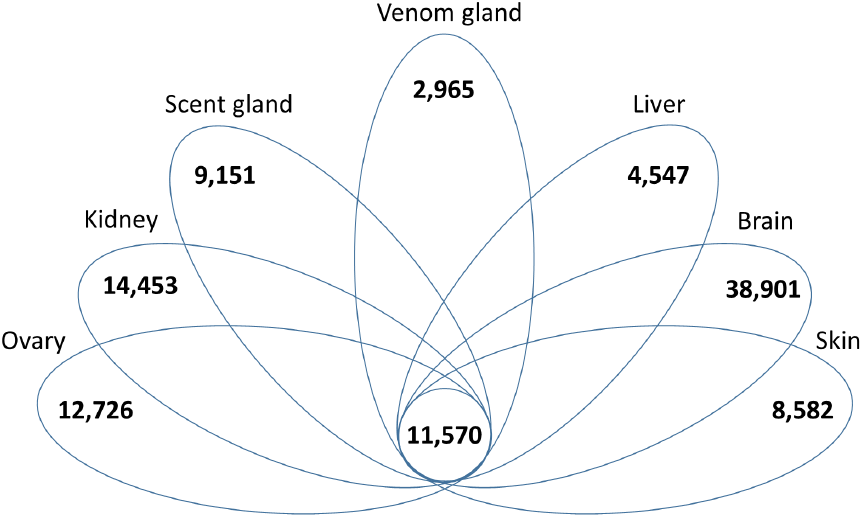
Tissue distribution of painted saw-scaled viper transcripts, determined by mapping sequencing reads derived from each tissue to a combined, all-tissue assembly with contig values of ≥1 FPKM (<u>F</u>ragments <u>P</u>er <u>K</u>ilobase of exon per <u>M</u>illion fragments mapped) taken as evidence for expression. Figures represent the number of unique transcripts expressed in each tissue, with the number of transcripts expressed in all 7 tissues indicated in the centre.

Venom gland samples for *Echis coloratus* were taken at different time points following manual venom extraction (“milking”); one sample approximately 16 hours post-milking, two samples at 24 hours and a final sample at 48 hours. We find 14,829 transcripts common to all 4 venom gland samples and suggest that these constitute the venom gland maintenance gene repertoire. However, we have also identified genes which were unique to each sample, with the 16 hour time point having 5,082 unique transcripts, the two 24 hour time points having 1,707 and 7,325 unique transcripts respectively, and the 48 hour time point having the highest number of unique transcripts with 12,535. Fishers exact tests revealed that the unique sequences expressed at 16 hours post-milking were mainly enriched for GO terms associated with transcription, such as “positive regulation of transcription from RNA polymerase II promoter”, “transcription initiation from RNA polymerase II promoter” and “RNA polymerase II transcription factor binding transcription factor activity” as well as “histone H3-K4 methylation” which is a known histone modification at the promoter of genes which are being actively transcribed (Liang et al. 2004; Santos-Rosa et al. 2002; Schneider et al. 2003; Schubeler et al. 2004). However, several categories relating to post-translational modification were also enriched, such as “protein SUMOylation” and “peptidyl-serine phosphorylation”. The GO terms “biosynthetic process” and “signaling” are also elevated compared to the other samples (Supplementary figure S10) as are “nucleic acid binding” and “protein binding” (Supplementary figure S11). At 24 hours, unique sequences are enriched for GO terms associated with translation, such as “translational initiation”, “translational elongation”, “structural constituent of ribosome” and “SRP dependent cotranslational protein targeting to membrane”. The mRNA surveillance pathway, “nuclear-transcribed mRNA catabolic process, nonsense-mediated decay” is also enriched. The 48 hour time point unique sequences had no significantly enriched GO terms when compared to either of the 24 hour samples, but were enriched for “catalytic activity” and “hydrolase activity” compared to the 16 hour sequences. A local BLAST survey of the venom gland transcriptomes revealed all previously characterized venom genes in *Echis coloratus* (Hargreaves et al. 2014b) were present by 16 hours post-milking.

### “Secretomics”

Between 3.72% and 10.64% of predicted open reading frames in the venom or salivary glands of the study species encoded a signal peptide and so are likely to be secreted (Supplementary table S2). Fishers exact tests show that the venom gland secretome of *E. coloratus* (based on pooled venom gland samples) is enriched for the GO terms “serine-type peptidase activity”, “peptidase activity acting on L-amino acid peptides”, “serine hydrolase activity” and “proteolysis” compared to the salivary gland secretomes of all non-venomous study species (Supplementary figures S1-S3). As viper venom contains primarily proteases including serine proteases, and is also known to contain L-amino acid oxidase, these results support an increased amount of these components being expressed in the venom gland compared to the salivary gland of non-venomous species. Interestingly the *E. coloratus* venom gland was also enriched for “serine-type peptidase activity” and “serine hydrolase activity” in comparison to the venom gland of the Eastern diamondback rattlesnake (*Crotalus adamanteus*), which may be indicative of interspecific variation in serine protease content in the venoms of these two species. When compared to the venom gland secretome of *E. pyramidum*, significant results were found only in the 16 and 48 hours post-milking samples, perhaps due to differences in the stages of the venom replenishment cycle between these samples and that of *E. pyramidum* venom gland (which was taken 24 hours post-milking), rather than any interspecific differences in gene expression. At 16 hours the venom gland secretome of *E. coloratus* was enriched for the GO terms “DNA binding”, “chromatin remodeling”, “nucleosome disassembly” and “transcription factor binding” compared to *E. pyramidum*. The GO categories “spliceosomal complex”, “protein polyubiquitination”, “protein transport”, “L-amino acid oxidase activity”, “serine-type endopeptidase inhibitor activity” and multiple categories for histone deacetylation were enriched in the 48 hours post-milking sample.

Sampling venom gland transcriptomes at different timepoints following milking allowed a comparison between venom gland secretomes at different stages of the venom replenishment cycle (Supplementary figures S13-S16). At 16 hours post-milking, ion binding and transferase activity appear to be elevated compared to other timepoints (Supplementary figure S14), and the GO terms “protein phosphorylation”, “transmembrane signaling receptor activity” and signal transducer activity” are significantly enriched compared to the remaining samples. The venom gland secretome 24 hours post-milking is significantly enriched for “protein ubiquitination”, “protein modification by small protein conjugation or removal”, “serine-type peptidase activity”, “zinc ion binding”, “peptidase activity” and “peptidase activity acting on L-amino acid peptides”. Peptidase activity appears to be considerably elevated in one 24 hour sample (Eco 7) but not in the other (Eco 6) (Supplementary figure S14) which may be suggestive of variation between individuals, but it may be more likely that this is a reflection of the difference in sequencing depth between these two samples (Supplementary table S1). After 48 hours several GO categories related to cellular components are elevated (Supplementary figure S15). We found this timepoint to be enriched for the highest number of GO categories, the majority of which were involved in histone deacetylation and chromatin modification. These include “chromatin modification”, “Histone deacetylase activity (H3-K9 specific)”, “protein deacetylase activity” and also “RNA splicing”. The prevalence of histone deacetylation at this timepoint suggests a reduction in the rate of gene transcription, as this process is known to cause transcriptional repression and gene silencing through the modification of higher-order chromatin structure (Ng and Bird 2000; Kouzarides 2007).

Finally, we also identified some variation in the number of secreted transcripts belonging to toxin gene families over time, suggesting that their expression may show temporal differences during the venom replenishment cycle (Supplementary figure S16). In general the number of secreted transcripts appears to reduce towards the 48 hour post-milking timepoint (as can be seen for metalloproteinase, C-type lectin, CRISP, VEGF-F and serine protease in Supplementary figure S16). This may be a result of the reduced rate of transcription indicated by the increase in histone deacetylation as mentioned previously. The notable exception to this is L-amino acid oxidase, with a single transcript being identified in the 16 hours and both 24 hours post-milking samples, but ten transcripts being identified at 48 hours post-milking. On further inspection we found that these transcripts in fact represented alternative splice variants which we previously characterized as *laao-b1* and *laao-b2* (Hargreaves et al. 2014b). The individual transcripts expressed at 16 and 24 hours post-milking encode *laao-b1*, which was found to be expressed in the venom gland of *E. coloratus* but also in the scent gland of royal python (*Python regius*), corn snake (*Pantherophis guttatus*) and rough green snake (*Opheodrys aestivus*) (i.e. all other snake species studied). The ten transcripts expressed 48 hours post-milking encode the variant *laao-b2* which we found to be venom gland-specific, and suggested it to be a putative venom component in *E. coloratus* due to its tissue specificity and elevated expression level (Hargreaves et al. 2014b). Whilst more sampling (including earlier and possibly later timepoints) is required to confirm temporal expression differences, and it is likely that not all potential venom gene sequences have been included unless they are present as a full length ORF in our dataset, it is certainly suggestive that venom gene expression may show temporal variation following venom expenditure.

When compared to other body tissues, the *E. coloratus* venom gland was found to be enriched for several GO categories including “peptidase activity”, “peptidase activity acting on L-amino acid peptides”, “serine-type peptidase activity” and “proteolysis” compared to brain; and for “protein modification by small protein conjugation or removal”, “protein ubiquitination” and “ligase activity” when compared to brain, ovary and scent gland. There were no GO terms significantly enriched in the venom gland compared to the remaining body tissues. A more complete comparison of GO terms between the venom gland and other body tissues of this species is represented by Supplementary figures S4-S6.

The venom gland of *E. pyramidum* was found to be enriched for “transforming growth factor beta receptor signaling pathway”, “negative regulation of macroautophagy”, “hedgehog receptor activity”, “serine-type endopeptidase activity” and “hormone secretion” compared to all salivary gland secretomes and the venom gland secretome of *E. coloratus*. When compared to the 48 hours post-milking sample of *E. coloratus*, it was also enriched for “protein glycosylation”, “metalloexopeptidase activity” and “cellular response to vascular endothelial growth factor stimulus”. In comparison to the venom gland secretomes of both *E. coloratus* and *E. pyramidum*, the venom gland secretome of the Eastern coral snake (*Micrurus fulvius*) was found to be enriched for “arachidonic acid secretion” (arachidonic acid is a fatty acid released from a phospholipid molecule following hydrolysis by a phospholipase A_2_ (Balsinde, Winstead, Dennis 2002). It is a precursor of the eicosanoids (such as prostaglandins) which can exert a diverse array of effects such as platelet aggregation, vasodilation and smooth muscle relaxation (Harizi et al. 2008)), “calcium-dependent phospholipase A_2_ activity”, “activation of phospholipase A_2_ activity” and “positive regulation of protein secretion”. This result is complementary to the finding that the venom gland transcriptome of this species consists predominantly of PLA_2_ transcripts (Margres et al. 2013) and that the toxicity of its venom is mainly due to PLA_2S_ (Vergara et al. 2014). No GO terms were found to be significantly enriched for the venom gland secretomes of king cobra or Eastern diamondback rattlesnake when compared to the venom gland secretome of either *Echis* species.

### Transcription factors

Our KEGG orthology (KO) analysis (Kanehisa et al. 2012) of the assembled transcriptomes identified between 255 and 358 transcription factors in the venom or salivary gland of our six study species, with lower numbers (between 40 and 143) in the smaller king cobra, Eastern coral snake and Eastern diamondback rattlesnake venom gland datasets (Table 1). We find 197 transcription factors to be conserved across the venom and salivary glands of our six study species, 77 of which are also found in all six additional *Echis coloratus* body tissues (ovary, liver, kidney, brain, cloacal scent gland and skin) and 71 of which are in five of the six tissues (Figure 3). We therefore suggest that these 148 widely-distributed transcription factors represent members of the basal transcription machinery common to all cells, although they may of course still play a role in the regulation of venom production. Interestingly, we find only two transcription factors which appear to unique to venom and salivary glands based on the available data; *SAM pointed domain-containing Ets transcription factor* (*Spdef*) and *Forkhead box A1/hepatocyte nuclear factor 3a* (*FoxA1/Hnf3a*) and of these *Spdef* is also found in the venom glands of the king cobra, Eastern coral snake and Eastern diamondback rattlesnake, whilst *FoxA1/Hnf3a* is also present in the Eastern diamondback venom gland. Two transcription factors were found only in the two *Echis* species; the homeodomain transcription factor *Ladybird* (*Lbx*) and *T-cell acute lymphocytic leukemia protein 1* (*Tal1*). Interestingly, we find no transcription factors common to all venomous species and the non-venomous colubrids (corn snake and rough green snake), suggesting that the loss of venom in these species may be the result of the loss of gene regulatory components. The king cobra accessory gland transcriptome appears to contain a much higher number of transcription factors than the venom gland (Table 1), with only 12 genes in common between the two. Only a single transcription factor appears to be unique to the venom gland of venomous snakes - the T-box transcription factor *Tbx3*, present in the venom gland of *Echis coloratus*, *Echis pyramidum*, Eastern diamondback rattlesnake, king cobra and Eastern coral snake, but not in the salivary gland of the leopard gecko, royal python, corn snake or rough green snake (transcripts encoding this gene are also found in four painted saw-scaled viper body tissues (kidney, ovary, scent gland and skin), but not king cobra pooled tissue or accessory gland).

**Table 1.** Numbers of transcription factors and components of signaling pathways found to be expressed in the venom (VG) or salivary gland (SAL). For completeness, we have also included data from Vonk et al. (2013) for the king cobra accessory gland (AG). There appears to be little difference between the venom gland of venomous snakes and the salivary glands of non-venomous species.

|  | Transcription<br>factors | Ca<sup>2+</sup><br>signalling | MAPK | NFκB | TGFβ | VEGF |
| --- | --- | --- | --- | --- | --- | --- |
| *Echis coloratus* (VG) | 307 | 74 | 139 | 67 | 51 | 31 |
| *Echis pyramidum* (VG) | 342 | 93 | 153 | 73 | 56 | 32 |
| Corn snake (SAL) | 308 | 60 | 137 | 58 | 51 | 30 |
| Rough green snake (SAL) | 255 | 59 | 125 | 49 | 44 | 31 |
| Royal python (SAL) | 278 | 65 | 138 | 55 | 48 | 31 |
| Leopard gecko (SAL) | 358 | 80 | 147 | 64 | 53 | 31 |
| Coral snake (VG) | 40 | 12 | 43 | 7 | 14 | 15 |
| Rattlesnake (VG) | 143 | 32 | 79 | 23 | 31 | 22 |
| King cobra (VG) | 40 | 21 | 26 | 7 | 14 | 11 |
| King cobra (AG) | 182 | 24 | 34 | 8 | 14 | 11 |

**Figure 3.**
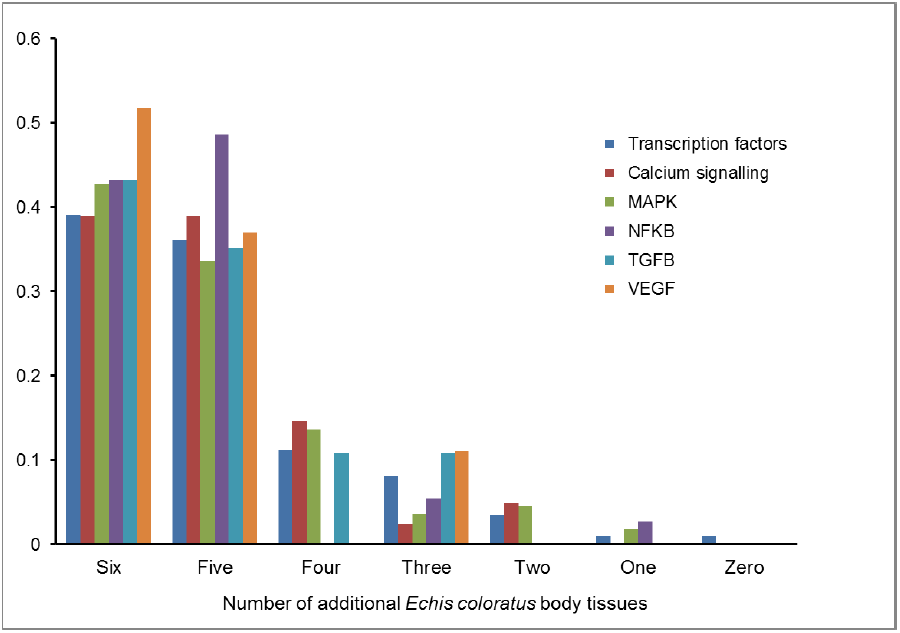
Proportions of transcription factors and members of signalling pathways found in the *Echis coloratus* venom gland that are also found in other body tissues.

It has previously been suggested that the transcription factors NF1, NFκB and AP-1 are involved in the regulation of snake venom production (Luna et al. 2009). Of the four known members of the nuclear factor 1 (Nf1) gene family (NFIA, NFIB, NFIC and NFIX (Kruse et al. 1991; Rupp et al. 1990)), we find only NFIA and NFIB to be expressed in the venom or salivary gland of all six study species. The NFκB family comprises five related transcription factors (RelA (p65), RelB, c-Rel, p50 (p105 precursor), p52 (p100 precursor)) that are able to form homo-or heterodimers via the shared Rel homology region (Hayden and Ghosh 2012; Napetschnig and Wu 2013). Of these, only RelA (p65), RelB, c-Rel encode a transactivation domain at their C-terminus and so are able to activate transcription as homodimers. The activity of NFκB is regulated via binding to the members of the IκB inhibitor protein family and sequestration of the resulting complex in the cytoplasm (Hayden and Ghosh 2012; Napetschnig and Wu 2013; Sen and Baltimore 1986). We find transcripts encoding RelA (p65), RelB, p50 (p105 precursor), p52 (p100 precursor) in the salivary or venom gland of all six study species and c-Rel in only Egyptian saw-scaled viper venom gland and corn snake salivary gland. We find transcripts encoding IκBα, IκBβ and IκBε inhibitors in all species. Activator protein 1 (AP-1) typically functions as a heterodimer of members of the Jun and Fos families of basic leucine zipper transcription factors (comprising c-Jun, JunB and JunD and c-Fos, FosB, FOS-like antigen 1 (FOSL1) and FOS-like antigen 2 (FOSL2)) (Curran and Franza Jr 1988; Karin et al. 1997). We find c-jun, junB and JunD and c-Fos and FOSL2 to be expressed in the venom or salivary gland of all six study species. Finally, *Sp1* binding sites have previously been identified upstream of the South American rattlesnake *crotamine* gene (Rádis-Baptista et al. 2003) and we find transcripts encoding this gene in the venom or salivary gland of all six study species.

### Signaling

The link between AP-1, NFκB and MAP kinase (ERK1/2) signalling has long been known (Hommes et al. 2003; Karin et al. 1997) and components of these pathways have previously been implicated in the regulation of venom production following milking in *Bothrops jararaca* (Kerchove et al. 2008; Luna et al. 2009), as has a role for calcium signalling (Kerchove et al. 2008), which is also known to be involved in salivary secretion (Melvin et al. 2005; Putney Jr 1986). The Sp1 transcription factor is known to interact with the Transforming Growth Factor-β (TGF-β) signalling pathway and members of the vascular endothelial growth factor (VEGF) gene family have been characterised from the venom gland of a diversity of reptile species (Aird et al. 2013; Francischetti et al. 2004; Margres et al. 2013; Yamazaki et al. 2009). Our analysis of members of these pathways in our salivary and venom gland transcriptomes (Table 1) revealed 110 transcripts involved in MAP kinase (ERK1/2) signalling that were conserved across the venom or salivary gland of our study species, 37 transcripts involved in NFκB signaling and an identical number involved in the TGF-β pathway, 41 transcripts involved in Ca^2+^ signaling and 27 in VEGF signaling. Total numbers of transcripts involved in each pathway were broadly similar across our study species, and slightly lower in the king cobra, Eastern diamondback and Eastern coral snake (Table 1). Interestingly, we found no evidence for venom-gland specific members of these signaling pathways, although RelB (involved in both the NFκB and MAPK pathways) and mitogen-activated protein kinase kinase kinase 2 (MAP3K2) from the MAPK pathway are expressed in only one of the six additional painted saw-scaled viper body tissues, the scent gland (Figure 3).

It has previously been suggested that α-and β-adrenoceptors (adrenergic receptors) may have a role in the activation of venom production following milking in *Bothrops jararaca* (Kerchove et al. 2008; Kerchove et al. 2004; Yamanouye et al. 2000) and a transcriptomic survey of the related *Bothrops alternatus* venom gland during venom synthesis three days post-milking suggested the presence of a conserved α_1D_ adrenoceptor (clone BACCGV4069B12, accession GW581578). However, our re-analysis of the *B. alternatus* data shows that this sequence in fact encodes a different G protein couple receptor, most likely a chemokine-like receptor. Nonetheless, we have identified several different adrenoceptors in our transcriptomic data (Table 2), although we find no evidence to support a conserved adrenoceptor in the venom gland of venomous snakes. We did not detect any adrenoceptor-like sequences in the venom glands of the king cobra, rattlesnake or coral snake and this, together with the lack of these sequences in previously published transcriptomes, suggests that a high sequencing depth is required to detect these transcripts. The apparent paucity of adrenoceptor transcripts in our data does not rule out their role in the regulation of venom production however, as it is possible (indeed likely) that these receptors are transcribed and translated prior to the synthesis of venom and therefore earlier than the time points studied by us and others.

**Table 2.** Presence of α and β adrenoceptor (adrenergic receptor) transcripts in reptile venom (VG) and salivary (SAL) glands. Eco, painted saw-scaled viper (*Echis coloratus*); Pgu, corn snake (*Pantherophis guttatus*); Oae, rough green snake (*Opheodrys aestivus*); royal python (Python regius); Ema, leopard gecko (*Eublepharis macularius*). Presence is denoted by a “+”, absence by a “-“.

|  | Species |  |  |  |  |
| --- | --- | --- | --- | --- | --- |
|  | Eco (VG) | Pgu (SAL) | Oae (SAL) | Pre (SAL) | Ema (SAL) |
| $\alpha 1a$ | + | + | - | - | + |
| $\alpha 1b$ | - | - | - | - | - |
| $\alpha 1c$ | - | - | - | - | - |
| $\alpha 1d$ | - | - | - | - | - |
| $\alpha 2a$ | - | - | - | + | + |
| $\alpha 2b$ | + | - | + | - | - |
| $\alpha 2c$ | - | - | - | - | - |
| $\alpha 2d$ | - | - | - | - | - |
| $\beta 1$ | + | - | - | + | + |
| $\beta 2$ | - | + | - | - | + |
| $\beta 4c$ | - | + | + | + | - |

### Comparative genomics

#### Genome assembly and completeness

Based on an estimated genome size of 1.3Gb for the painted saw-scaled viper (derived from the haploid genome size of 1.27Gb for the related *Echis carinatus* (Desmet 1981)) we have sequenced this genome to a depth of roughly 89.2x according to the Lander/Waterman equation (number of reads x read length/genome size (Lander and Waterman 1988); full assembly statistics are given in Supplementary table S7). Our assembly has a contig N50 of 3.86kb, roughly similar to the 3.98kb of the only other venomous snake genome sequence currently available (the king cobra (Vonk et al. 2013)), although our scaffold N50 of 5.58kb is considerably lower than theirs (226kb), most likely reflecting the absence of mate-pair library data in our assembly. Our assembly contains 1,097 gene-sized scaffolds (here gene-sized is defined as being ≥25kb in length (Bradnam et al. 2013)). Analysis of the conserved eukaryotic gene set using the Core Eukaryotic Genes Mapping Approach (CEGMA (Parra et al. 2007)) identified 81 full length genes out of the 248 highly conserved core eukaryotic genes (CEGs), suggesting that our genome assembly is approximately 33% complete (compared to 56% and 49% for the Burmese python and king cobra respectively). However, we identified 195 partial CEGs, giving a completeness figure nearer 79% (96% for Burmese python and 92% for king cobra). We should therefore expect to find partial matches to most of our genes of interest in our genome assembly, although for prediction of transcription factor binding sites and comparative analyses of promoter regions it is the region immediately upstream of the first exon or transcription start site (where known) that is most interesting.

#### Transcription factor binding sites analysis

We interrogated our genome sequence for genes known to encode venom toxins in *E. coloratus* and compared the upstream regions of these genes (where data were available) with other members of the same gene family within this species and to related genes in other species using previously published data.

### Snake venom metalloproteinases

Our *Echis coloratus* genome assembly contains 15 scaffolds encoding the first exon of one of the many snake venom metalloproteinase gene variants (SVMP, a group of metalloproteinases thought to be most closely related to the A Disintegrin And Metalloproteinase family of peptidases (Casewell 2012; Jia et al. 1996)). Of these, eight had typically only a few hundred base pairs of sequence upstream of the start codon and a longer sequence had a string of N’s adjacent to the start codon. These 9 sequences were eliminated form subsequent analyses. The remaining seven contigs had ≥1000bp of sequence upstream of the start codon and provided 2,533bp of aligned sequence (based on the length of the shortest aligned sequence) for transcription factor binding site analysis. Although present in all sequences, we find little conservation of predicted binding sites for candidate transcription factors such as AP-1, AP-2, Sp1, NF1, NFκB, Tbx3, Hnf3a across most of these regions, although a roughly 500bp region immediately upstream of the start codon does contain sites conserved across several (NF1 and NFκB) or all (Nkx2.5, Tef1, Barbie, ETS) sequences (Figure 4), suggesting that this proximal region may be most important in controlling gene expression.

**Figure 4.**
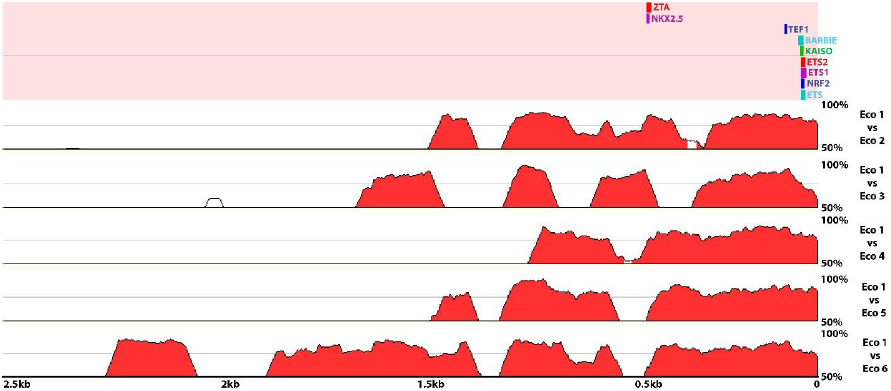
Transcription factor binding site analysis of 2,533bp of upstream sequence of six *Echis coloratus* (Eco) snake venom metalloproteinase (SVMP) genes. Position 0 denotes the first base of the start codon. The locations and approximate length of the nine transcription factor binding sites found to be conserved across all six sequences are shown in the top panel and pairwise sequence similarity plots are shown in the bottom panels, with peaks over 75% conserved colored red. Conserved transcription factor binding sites and the highest levels of sequence conservation are seemingly restricted to a region proximal to the transcription start site.

### Serine proteases

There are eight scaffolds in our assembly that encode the first exon of a serine protease gene and we were able to generate an alignment of at least 1500bp for six of these. We find evidence for a number of conserved transcription factor binding sites within approximately 500bp of the start codon of all of these genes (Figure 5), although not for any of the candidate transcription factors.

**Figure 5.**
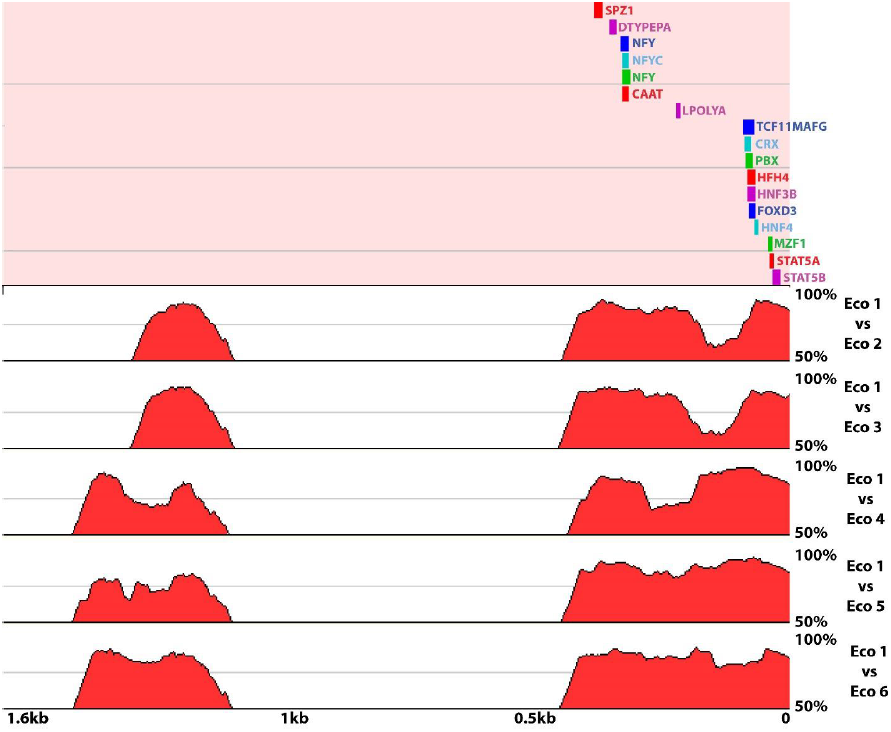
Transcription factor binding site analysis of 1,624bp of upstream sequence of six *Echis coloratus* (Eco) serine protease genes. Position 0 denotes the first base of the start codon. The locations of the seventeen transcription factor binding sites found to be conserved across all six sequences are shown in the top panel and pairwise sequence similarity plots are shown in the bottom panels, with peaks over 75% conserved colored red. As for SVMP genes (Figure 4), conserved transcription factor binding sites are seemingly restricted to a region proximal to the transcription start site.

### C-type lectins

Our *E. coloratus* genome assembly contains thirteen scaffolds that encode the first exon of a C-type lectin gene. Six of these scaffolds contain less than 150bp of sequence upstream of the start codon and one contained a long string of N’s immediately adjacent to the start and these seven sequences were therefore excluded from further analysis. The remaining six sequences provided 402bp of aligned sequence and although each contains putative binding sites for candidate transcription factors there are no conserved sites shared by all loci. However, analysis of mRNA sequences reveals that transcripts expressed in the venom gland have a 12 nucleotide insertion in their 5’ UTR which is not present in the C-type lectin transcript expressed in the scent gland of this species. We found no transcription factor binding sites within this region, although the first 3 nucleotides of the insertion have led to the formation of a binding site for the transcriptional activator MYB.

### Other venom genes

We find two scaffolds in our assembly that encode Group IIA Phospholipase A2(PLA_2_) genes and these provided 292bp of aligned sequence. The upstream regions of these two genes are highly identical and contained 56 conserved transcription factor binding sites, including sites for Spz1, Tbx, NF1, Mzf1, Sp1, HFH4. Two scaffolds in our assembly encode CRISP genes and these provided 2.2kb of upstream sequence for analysis, resulting in the identification of 136 conserved transcription factor binding sites. It seems likely that the level of sequence conservation between both PLA_2_ and CRISP paralogs is too high for the accurate prediction of putative transcription factor binding sites in this species. Finally, our *E. coloratus* genome sequence contains an 8.3kb scaffold that encodes the full coding sequence of the *vascular endothelial growth factor f* (*vegf-f*) gene (a viper-specific paralog which appears to be venom gland-specific (Hargreaves et al. 2014b)), including 2.4kb of sequence upstream of the transcription start site. This upstream region contains putative transcription factor binding sites for AP-1, AP-2, Tbx and NFκB (among others), although a more detailed comparison is not currently possibly due to a lack of sequence for the upstream regions of other vegf genes in this species or others.

### Multi-species comparisons

A 25,026bp cosmid sequence encoding a cluster of three functional PLA_2_ genes and two non-functional pseudogenes has previously been published for the Okinawan habu (*Protobothrops flavoviridis*) (Ikeda et al. 2010). We searched the upstream regions of each of the functional genes (defined as the region upstream of the start (ATG) codon to the stop codon of the preceding gene or pseudogene, comprising 1,829bp upstream of PfPLA-2; 4,045bp upstream of PfPLA-5 and 5,444bp upstream of PfPLA-4) and identified 168 putative transcription factor binding sites conserved across the three sequences (an unsurprising finding given the high level of sequence identify between the three regions). These included conserved sites for candidate transcription factors such as AP-1, AP-2, Sp1, NFκB, Tbx3 and Hnf3a and, most interestingly, a conserved Tef1 binding site, similar to that seen in *E. coloratus* SVMP loci (Figure 4). Comparison of these *Protobothrops* sequences to those from our *E. coloratus* genome identified 9 conserved transcription factor binding sites (Figure 6), all located within a few hundred base pairs of the start codon.

**Figure 6.**
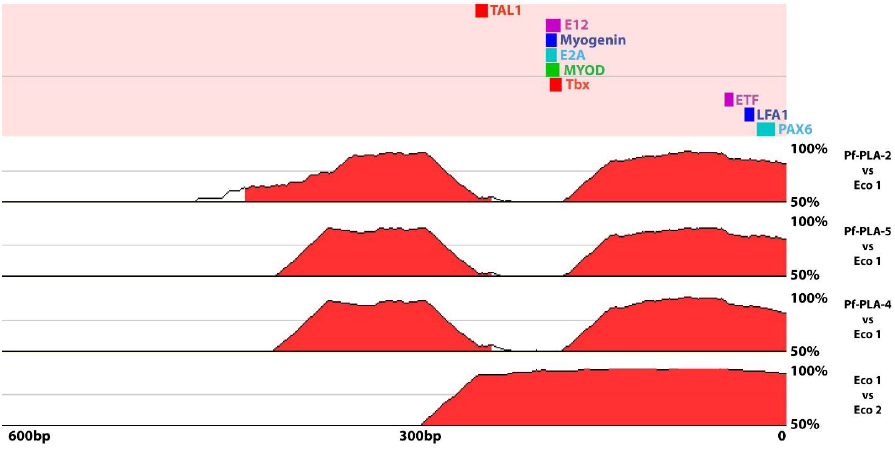
Transcription factor binding site analysis of the upstream regions of *Echis coloratus* (Eco) and *Protobothrops flavoviridis* (Pfl) phospholipase A_2_ (PLA_2_) genes. Position 0 denotes the first base of the start codon. The locations of the nine transcription factor binding sites found to be conserved across all sequences are shown in the top panel and pairwise sequence similarity plots are shown in the bottom panels, with peaks over 75% conserved colored red.

The full coding regions and approximately 300bp of upstream sequence for 19 PLA_2_ genes from four North American rattlesnakes (*Sistrurus catenatus edwardsi*, *S. c. termgeminus*, *S. c. catenatus* and *Sistrurus miliarius barbouri*) have previously been published (Gibbs and Rossiter 2008). The upstream regions of these genes are highly similar, although there has been a 42bp insertion in one of the *S. c. edwardsi* sequences (edw4, accession EU369751) which has been annotated as a pseudogene. We compared the remaining 18 genes to the upstream regions of the two *E. coloratus* and three *Protobothrops* PLA_2_ sequences and identified three conserved transcription factor binding sites, for E2A, myogenin and Tbx.

Although the level of sequence conservation between the *E. coloratus* CRISP paralogs was too high to accurately predict conserved transcription factor binding sites, a comparison incorporating an upstream region of a king cobra CRISP gene (on scaffold 7136, accession AZIM01007132) resulted in a much smaller list of just 35 conserved TF binding sites, all located within 2kb of the start of the gene (Figure 7).

**Figure 7.**
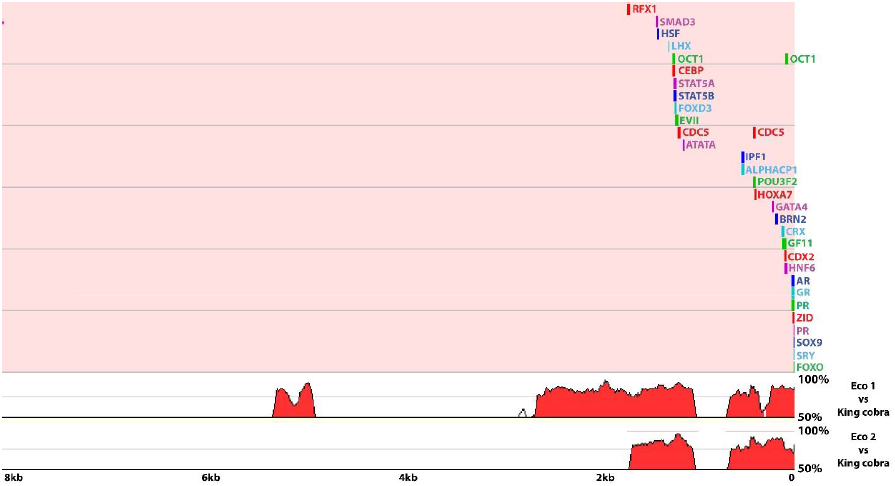
Transcription factor binding site analysis of 7,272bp of upstream sequence of *Echis coloratus* (Eco) and king cobra (*Ophiophagus hannah*) cysteine-rich secretory protein (CRISP) genes. Position 0 denotes the first base of the start codon. The locations of the 33 transcription factor binding sites found to be conserved across all sequences are shown in the top panel and pairwise sequence similarity plots are shown in the bottom panels, with peaks over 75% conserved colored red.

We examined the available genomic scaffolds for nerve growth factor (*ngf*, accessions AZIM01002615 and AZIM01012844) from the king cobra (*Ophiophagus hannah*) (Vonk et al. 2013) which has previously been suggested to have undergone a gene duplication within the Elapid lineage (Hargreaves et al. 2014a; Hargreaves et al. 2014b; Sunagar et al. 2013). Based upon phylogenetic analysis (Hargreaves et al. 2014a) we call these *ngfa* and *ngfb*, with *ngfb* being the putatively toxic version due to its selective expression in the venom gland. We also extracted *ngfa* sequences for the painted saw-scaled viper, the corn snake, the Burmese python (Castoe et al. 2013) (accession KE954116,) and the boa constrictor (Bradnam et al. 2013) (scaffold SNAKE00002822) to aid in the identification of unique transcription factor binding sites. All six upstream regions were aligned to give 528bp of sequence upstream from the start codon for transcription factor binding site analysis. The total number of transcription factor binding sites was found to vary between species with 98 and 88 in king cobra *ngfa* and *ngfb* respectively, 163 in the painted saw-scaled viper, 101 in corn snake, 96 in Burmese python and 88 in boa constrictor. Just twelve transcription factor binding sites were found to be conserved between all species (CBF, USF, EBOX, BEL1, CMAF, PAX9, NRSE, NRSF and 2 sites for both YY1 and MYCMAX) (Figure 8). In the king cobra, *ngfa* and *ngfb* have 57 transcription factor binding sites in common. 28 additional sites were found to be conserved between king cobra *ngfa* and *ngfa* from other species, but are missing from king cobra *ngfb*. We found 13 binding sites unique to king cobra *ngfa*, most noticeably 3 binding sites for SMADs which are not present in any other *ngfa* upstream region from other species, or in *ngfb*. The putatively toxic *ngfb* has 70 sites conserved between itself and *ngfa* in either king cobra or at least one other snake species. We found 13 novel transcription factor binding sites in *ngfb* including one for hepatocyte nuclear factor (HNF4), one site for TATA-binding protein (TBP) and one site for transcription factor IIA (TFIIA) and there has also been an 8 nucleotide deletion 409bp upstream from the start codon.

**Figure 8.**
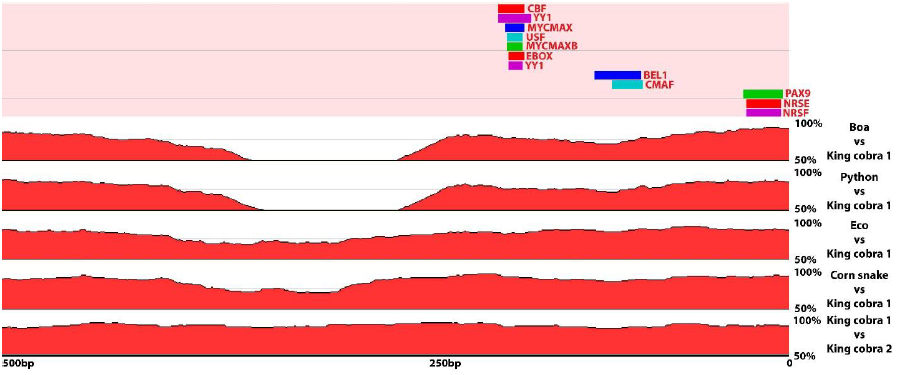
Transcription factor binding site analysis of 528bp of upstream sequence of *Echis coloratus*, king cobra (*Ophiophagus hannah*), corn snake (*Pantherophis guttatus*), Burmese python (*Python molurus bivittatus*) and Boa constrictor (*Boa constrictor constrictor*) nerve growth factor (NGF) genes. Position 0 denotes the first base of the start codon. The locations of the 12 transcription factor binding sites found to be conserved across all sequences are shown in the top panel and pairwise sequence similarity plots are shown in the bottom panels, with peaks over 75% conserved colored red.

An insertion upstream of the transcription start site of the *coagulation factor X* gene in the rough-scaled snake (*Tropidechis carinatus*) and Eastern brown snake (*Pseudonaja textilis*) has been claimed to be responsible for the increased expression level of a venom gland-specific paralog following gene duplication (Kwong et al. 2007; Reza et al. 2007), and accordingly we find no evidence for either gene duplication or this insertion in *E. coloratus*, corn snake, king cobra or Burmese python. The upstream regions of the duplicated factor X genes (pseutarin C and trocarin D (Rao et al. 2004; Reza et al. 2007)) are extremely similar and possess 147 conserved transcription factor binding sites.

## DISCUSSION

Our comparative transcriptomic and genomic analyses provides evidence that a diversity of gene regulatory networks may be involved in the transcriptional regulation of genes encoding snake venom toxins. Analyses of different time points during venom synthesis following manual extraction of venom reveals an apparent shift from a focus on transcription to translation between 16 and 24 hours post milking, indicating that activation of genes encoding venom toxins is a relatively rapid event. We identified only a single transcription factor (*Tbx3*) that appears to be unique to the venom glands of venomous snakes and absent from the salivary glands of non-venomous species. Whilst more work is needed to establish the distribution of this gene in additional species, its presence in the venom gland is intriguing, since TBX3 is a known transcriptional repressor and is known to interact with diverse gene regulatory networks in multiple tissues in a context-dependent manner (Washkowitz et al. 2012). If TBX3 is carrying out a similar repressive function in the venom gland, then it is possible that the initiation of venom production following expenditure is facilitated by the removal of transcriptional repression (i.e. negative regulation), rather than by transcriptional activation. A similar situation is known to occur during embryonic development, where RNA polymerases are ‘paused’ on gene promoters and in this way offer a rate-limiting mechanism for transcription (Core and Lis 2008; Krumm et al. 1995). Such a system also enables rapid initiation of gene expression, as the transcriptional machinery is already assembled and in place on the gene itself (Margaritis and Holstege 2008). A gene regulatory mechanism such as this obviously has important implications for the rapid initiation of venom replenishment, although the lack of conserved TBX binding sites in the upstream regions of venom genes would suggest that TBX3 is acting higher up in the venom gene regulatory network.

The presence of NFκB and its inhibitors in the venom glands of all species, together with the previous identification of a role for these transcription factors in the regulation of venom production (Luna et al. 2009) also supports a mechanism of rapid initiation of venom replenishment following expenditure, as NFκB dimers are held inactive in the cytoplasm via association with inhibitor proteins and can be rapidly activated by the removal and degradation of these inhibitors (Karin 1999).

If different gene families are regulated by distinct gene regulatory networks (or sub-components of the same network) then we might expect to see differences in the temporal expression of these genes. Our results indicate that this may be the case, with evidence for the expression of some gene families declining quite rapidly after 16 hours, and some increasing quite dramatically (such as *laao-b2*) after 48 hours. Not only is this suggestive of a number of gene regulatory networks being involved, it also has implications for the importance of respective venom toxins, with those essential to the functional efficacy of the venom being expressed and replaced almost immediately following expenditure.

Our draft painted saw-scaled viper (*Echis coloratus*) genome enabled us to carry out comparative analyses of the upstream regions of the gene families that make the most substantial contribution to the venom of this and closely related species (SVMPs, Phospholipase A_2S_, C-type lectins and Serine Proteases) (Casewell et al. 2009; Casewell et al. 2014; Wagstaff et al. 2009). These analyses identified a number of conserved predicted transcription factor binding sites between members of gene families, but not between members of different gene families, supporting our proposal that multiple gene regulatory networks may be acting within the snake venom gland, each working to activate the expression of typically one gene family. This situation is likely a reflection of multiple restriction events at different times during the evolution of venomous snakes, where a gene encoding a salivary protein has been duplicated and the expression of one of the copies restricted to the venom gland (Hargreaves et al. 2014a). It is also likely that the different transcriptional networks acting on the different gene families contribute to observed differences in venom composition within and between species (Casewell et al. 2014; Chippaux et al. 1991). In all cases, the conserved transcription factor binding sites were located within a few hundred base pairs of the start of the gene and this, together with the observed short intergenic distances in the previously characterized Okinawan habu (*Protobothrops flavoviridis*) PLA_2_ ene cluster (Ikeda et al. 2010), supports our initial hypothesis and explains the apparent ease with which functional copies of existing venom genes are produced via gene duplication (Kordiš and Gubenšek 2000; Wong and Belov 2012). It remains to be seen however if the short regulatory regions of these genes are an ancestral condition, in which case they may be considered to be pre-adapted or exapted (Gould and Vrba 1982) to repeated gene duplication, or if only partial regulatory regions were initially duplicated, in which case these genes may have been subfunctionalised and restricted to the venom gland by default.

Finally, our comparative transcriptomic analysis of the *E. coloratus* venom gland with other body tissues in this species, and with the venom and salivary glands of other species, has resulted in some intriguing findings. Not only does the venom gland express the lowest number of unique transcripts of any of the seven tissues we have investigated, but it also does not appear to be particularly outstanding with respect to the number and type of secreted products. Whilst snake venom itself may represent an evolutionary innovation, with extensive intra-and inter-specific variation in venom composition resulting from a range of genomic, transcriptional, translational and post-translational mechanisms, the venom gland itself may be just another tissue.

## MATERIALS AND METHODS

All research involving animals was carried out in accordance with institutional and national guidelines and was approved by the Bangor University Ethical Review Committee.

### RNA-Seq

RNA-Seq-derived transcriptomes for painted saw-scaled viper, corn snake, rough green snake, royal python and leopard gecko were carried out as described previously (Hargreaves et al. 2014a; Hargreaves et al. 2014b) and we use the general term ‘salivary gland’ for simplicity, to encompass the oral glands of the leopard gecko and rictal glands and Duvernoy’s gland of the royal python, corn snake and rough green snake and do not imply any homology to mammalian salivary glands. Egyptian saw-scaled viper venom gland data was derived from an adult female specimen sacrificed 24 hours following manual venom extraction using the same methodology. Briefly, total RNA was extracted using the RNeasy mini kit (Qiagen) with on-column DNase digestion and mRNA sequencing libraries were subsequently prepared using the TruSeq RNA sample preparation kit (Illumina) with a selected insert size of 200-500bp. Libraries were then sequenced on the Illumina HiSeq2000 or HiSeq2500 platform using 100bp paired-end reads. The resulting data were assembled using Trinity (Grabherr et al. 2011; Haas et al. 2013) and putative open reading frames (ORFs) extracted using longorf.pl (available online at http://bip.weizmann.ac.il/adp/bioperl/bioSeq/longorf) using the strict option (where ORFs must start with an ATG). Signal peptides were predicted using SignalP (Petersen et al. 2011) (version v4.1). Transcript annotation and assignment of gene ontology (GO) terms was performed using BLAST2GO (Conesa et al. 2005), the KEGG Automatic Annotation Server (KAAS) (Moriya et al. 2007) and by local BLAST using BLAST+ (Camacho et al. 2009) (version v2.2.27). Enrichment analyses using Fisher exact tests were implemented using BLAST2GO. Tissue distribution of transcripts was determined through the creation of a global all tissue assembly and read mapping/transcript abundance estimation using RSEM (Li and Dewey 2011) as a downstream application of Trinity (version r2013-02-25), with an FPKM (<u>F</u>ragments <u>P</u>er <u>K</u>ilobase of exon per <u>M</u>illion mapped reads) value of ≥1 taken as confirmation of expression.

### Genome sequencing

A draft genome sequence for the painted saw-scaled viper was generated from a single adult female (heterogametic ZW) specimen originating from Israel. Genomic DNA was extracted from muscle tissue using phenol/chloroform. Two genomic libraries were prepared using the Illumina Truseq sample preparation kit with insert sizes of 300bp and 600bp by the GenePool at the University of Edinburgh (http://genepool.bio.ed.ac.uk/). These were pooled and sequenced on one lane of the Illumina HiSeq 2000 sequencing platform, producing a total of 579,767,826 100bp paired-end reads (see table S7 for full sequencing metrics). The resulting reads were assembled using the short read *de novo* assembler ABySS (Simpson et al. 2009) with a k-mer size of 60. The resulting assembly was assessed for completeness using the Core Eukaryotic Genes Mapping Approach (CEGMA) (Parra et al. 2007) using the pipeline version v2.4.010312. Available genome sequences for the Burmese python (Castoe et al. 2013) and the king cobra (Vonk et al. 2013) were also incorporated into the CEGMA analysis to allow comparison between assemblies.

### Transcription factor binding sites analysis

Putative transcription factor binding sites were annotated using MultiTF, implemented through Mulan (Ovcharenko et al. 2005) with a 100bp conserved region length and 70% conservation limit. Upstream region sequences were first aligned using ClustalW (Larkin et al. 2007) and then trimmed to leave only contiguous sequence leading up to the start (ATG) codon. Sequences were then surveyed for all available evolutionarily conserved transcription factor binding sites, specifically for transcription factors previously implicated to have a role in the regulation of snake venom production, including NFκB and AP-1, as well as those predicted by the current study, such as Tbx3. The genome assembly for the king cobra was downloaded from GenBank under the accession AZIM00000000.1. The assembly for the Burmese python genome (v2.0) was downloaded from the snake genomics website (http://www.snakegenomics.org/SnakeGenomics/Available_Genomes.html) and this assembly is also available on GenBank under the accession AEQU00000000.2. All genome assemblies of the Boa constrictor generated for Assemblathon 2 (Bradnam et al. 2013) are available for download from GigaDB (http://gigadb.org/dataset/view/id/100060). We used the assembly “snake_1C_scaffolds” which was assembled by the BCM-HGSC team (for more details on the assembly method used see Additional file 3 in (Bradnam et al. 2013)).

## Acknowledgments

The authors wish to thank R. Morgan, A. Barlow and C. Wüster for technical assistance. We would also like to acknowledge the always enthusiastic help and support of the late Ashley Tweedale. We are very grateful to the staff of High Performance Computing (HPC) Wales for enabling and supporting our access to their systems, and thank Richard Storey and Daniel Struthers of PetGen Ltd. for their partnership during this project.

### FINANCIAL DISCLOSURE

This research was partially supported by a Royal Society Research Grant awarded to JFM and Wellcome Trust funding to DWL (grant number 098051). JFM, MJH and MTS are supported by the Biosciences, Environment and Agriculture Alliance (BEAA) between Bangor University and Aberystwyth University and ADH is funded by a Bangor University 125th Anniversary Studentship.

### COMPETING INTEREST

The authors have no competing interests.

### DATA ACCESS

Transcriptome reads were deposited in the European Nucleotide Archive (ENA) database under accession #ERP001222 and in the NCBI Sequence Read Archive (SRA) under the study accessions #SRP042007 and #SRP043460. Sequencing reads for the *Echis coloratus* genome were deposited in the SRA under study accession #SRP043211.

